# Identifying tagging SNPs for African specific genetic variation from the African Diaspora Genome

**DOI:** 10.1101/112235

**Authors:** Henry Richard Johnston, Yi-Juan Hu, Jingjing Gao, Timoty D. O’Connor, Goncalo Abecasis, Genevieve L Wojcik, Christopher R. Gignoux, Pierre-Antoine Gourraud, Antoine Lizee, Mark Hansen, Rob Genuario, Dave Bullis, Cindy Lawley, Eimear E. Kenny, Carlos Bustamante, Terri H. Beaty, Rasika A. Mathias, Kathleen C. Barnes, Zhaohui Steve Qin, CAAPA Consortium

**Affiliations:** Department of Biostatistics and Bioinformatics, Emory University, Atlanta, GA; Data and Statistical Sciences, AbbVie, North Chicago, IL; Institute for Genome Sciences, University of Maryland School of Medicine, Baltimore, MD; Program in Personalized and Genomic Medicine, University of Maryland School of Medicine, Baltimore, MD; Department of Medicine, University of Maryland School of Medicine, Baltimore, MD; Department of Biostatistics, SPH II, University of Michigan, Ann Arbor, MI; Department of Genetics, Stanford University School of Medicine, Stanford, CA; Department of Neurology, University of California, San Francisco, San Francisco, CA; Illumina, Inc., San Diego, CA; Department of Genetics and Genomics, Icahn School of Medicine at Mount Sinai, New York, NY; Department of Epidemiology, Bloomberg School of Public Health, JHU, Baltimore, MD; Department of Medicine, Johns Hopkins University, Baltimore, MD

**Author notes:** Correspondence and requests for materials should be addressed to Kathleen C. Barnes. A full list of consortium members appears at the end of the Supplement (CAAPA Consortium).

## Abstract

A primary goal of The **<u>C</u>**onsortium on **<u>A</u>**sthma among **<u>A</u>**frican-ancestry **<u>P</u>**opulations in the **<u>A</u>**mericas (CAAPA) is to develop an ‘African Diaspora Power Chip’ (ADPC), a genotyping array consisting of tagging SNPs, useful in comprehensively identifying African specific genetic variation. This array is designed based on the novel variation identified in 642 CAAPA samples of African ancestry with high coverage whole genome sequence data (~30x depth). This novel variation extends the pattern of variation catalogued in the 1000 Genomes and Exome Sequencing Projects to a spectrum of populations representing the wide range of West African genomic diversity. These individuals from CAAPA also comprise a large swath of the African Diaspora population and incorporate historical genetic diversity covering nearly the entire Atlantic coast of the Americas. Here we show the results of designing and producing such a microchip array. This novel array covers African specific variation far better than other commercially available arrays, and will enable better GWAS analyses for researchers with individuals of African descent in their study populations. A recent study^1^ cataloging variation in continental African populations suggests this type of African-specific genotyping array is both necessary and valuable for facilitating large-scale GWAS in populations of African ancestry.

## Introduction

The design of the African Diaspora Power Chip (ADPC) was a primary goal as part of the NIH-supported **<u>C</u>**onsortium on **<u>A</u>**sthma among **<u>A</u>**frican-ancestry **<u>P</u>**opulations in the Americas (CAAPA). Because of the overall poor coverage for African specific-variants on commercially available GWAS arrays, amongst other difficulties, relatively few GWAS have been performed in populations of African descent^2^, in part because they were underpowered to identify association with genes controlling risk for complex disease^3^. Previous GWAS studies in populations of African descent may have missed critical association signals because the single nucleotide polymorphisms (SNPs) genotyped on existing commercial arrays were selected for being informative among individuals of European ancestry, and generally do a poor job of tagging haplotypes and variants in individuals of non-European ancestry. This stems from the fact that the frequency of SNPs on currently available arrays is not well matched to the frequency of untagged variants in non-European populations. Essentially, the variant spectrum on current SNP arrays is flat, ensuring common genetic variants are well tagged, but making it difficult to tag low frequency SNPs (even though they may be highly polymorphic in non-European populations). This missing genetic variation in non-Europeans, however, consists largely of low frequency and rare variants, which will always be poorly covered by tagging SNPs with higher minor allele frequency (MAF). By building a large catalog of novel African-specific genetic variants, and then designing an array to tag as many of these as possible, we provide researchers with a significantly improved tool for hunting genes associated with diseases in populations of African ancestry, including admixed populations.

Ongoing work in the CAAPA consortium has included coverage analysis of the novel variation identified by CAAPA sequencing. This analysis has shown that only 69% of common SNP variants and 41% of low-frequency SNP variants identified by CAAPA can be tagged by traditional GWAS arrays (at r^2^ >= 0.8), such as the Illumina HumanOmni5 (which contains about five million SNPs), [Mathias et al. 2016]. Ha, et al.^4^ suggested much lower coverage levels, with the OmniExpress chip (containing about 770,000 SNPs) effectively covering only 8% of known variation within the YRI genome, while the much larger Omni 2.5 (containing about 2.5 million SNPs) still only covers 20% of known YRI SNPs based on their analysis. In contrast, variants of European (CEU) ancestry are 21% covered by the OmniExpress and 44% covered by the Omni 2.5. These are large differences in genomic coverage, and are supported by other studies with similarly pessimistic estimates of effective coverage among non-European populations^5-7^. Even the primary manufacturer of the Omni series genotyping chips, Illumina, refers to low coverage levels for their currently available commercial GWAS arrays in non-European populations. [Table 1] It is important to note that there is no standardized definition for efficient ‘coverage,’ as each method uses a different set of SNPs to assess coverage levels. Regardless of the method used, however, contemporary commercially available arrays do a poor job of tagging common haplotypes or ‘covering’ all genetic variation in non-European or admixed populations.

**Table 1.** Illumina projected coverage of African variants on several commercially available GWAS arrays

| Table 1 | MAF Category |  |  |  |
| --- | --- | --- | --- | --- |
|  | <0.010<br>(non-singletons) | >0.010 | >0.025 | >0.050 |
| HumanExomev1-2 | 0.031 | 0.032 | 0.034 | 0.035 |
| HumanCore-12v1-0 | 0.127 | 0.152 | 0.203 | 0.256 |
| HumanCoreExome-12v1-0 | 0.148 | 0.172 | 0.221 | 0.271 |
| OmniExpress-12v1-1 | 0.21 | 0.249 | 0.326 | 0.395 |
| OmniExpressExome-8v1-1 | 0.226 | 0.263 | 0.337 | 0.403 |

Usage of the most recent imputation panels can significantly improve the coverage of African variants, but this practice is still hamstrung by the lack of low-frequency variants on genotyping arrays (MAF < 5%). Imputation of low-frequency variants is most efficient and accurate when the SNP to be imputed has a similar minor allele frequency as the genotyped SNP, so a relative lack of low frequency variants on an array can render imputation of similar frequency variants difficult [Marchini and Howie, 2010].

To address this shortcoming, the ADPC was designed using the whole-genome sequencing results on 642 CAAPA samples, including 328 African Americans, 125 African Caribbean subjects, 164 African ancestry individuals with some Latino ancestry, and 25 individuals from Nigeria. The whole genomes of these individuals were sequenced using the Illumina HiSeq 2000. A total of 47.9 million biallelic SNPs were identified in these CAAPA samples. Of those, 15.6 million variants have a MAF greater than or equal to 1%.

To create an affordable array for large-scale chip-based studies, 700,000 variants was the maximum size of the array. A MAF of 1% was chosen as the preliminary cutoff to limit the initial pool of variants to be tagged. In addition, using 1% as the MAF cut-off eliminated concerns about potential false positive variant calls for rare variants derived from the sequence data^8^. Additionally, the ADPC is designed to be used in conjunction with the OmniExpress array, a low-cost GWAS array popular among researchers, leveraging the high MAF coverage available from OmniExpress and freeing the ADPC to focus on low frequency variants.

To narrow the pool further, SNPs with poor Illumina design scores were removed from consideration as potential tag SNPs. MaCH^9^ and minimac^10^ were used to determine which CAAPA variants were well-imputable based on the 1000 Genomes Phase I African Reference Panel and variants from the OmniExpress array. Those SNPs that could be imputed well (r^2^ > 0.8) were removed from the pool of SNPs needing to be tagged.

Fugue^11^ was then used to determine the pairwise linkage disequilibrium (LD) between each pair of SNPs in the remaining set of SNPs. These LD estimates were used by FESTA^12^, a TagSNP selection program, to select SNPs using a rather strict r^2^ threshold of 0.8. A total of 1,004,268 TagSNPs were selected. Among them, 4,000 were removed because they were too similar in their probe design to function well on the array. Limited by the capacity of the array, only TagSNPs with an MAF >= 1.6% were retained for inclusion on the array. Additional content was then added. This includes SNPs previously found to be associated with African-specific diseases but not previously selected as TagSNPs and approximately 600 additional SNPs in the human leukocyte antigen (HLA) region. HLA SNPs added to the array are relevant for HLA imputation and analyses of diseases or phenotypes related to immunity. These SNPs represent a pruned selection of SNPs with high tagging power, directly through LD, as well as SNPs preferentially selected by existing SNP-based HLA imputation algorithms^13^. Finally, a “GWAS fingerprint” that includes 274 markers, identical to that used on the Illumina HumanExome array, was added to enable researchers to ensure accurate sample labeling and analysis, while facilitating sample tracking across experiments. In final count, 627,998 variants were included on the ADPC array. [Figure 1]

**Figure 1.**
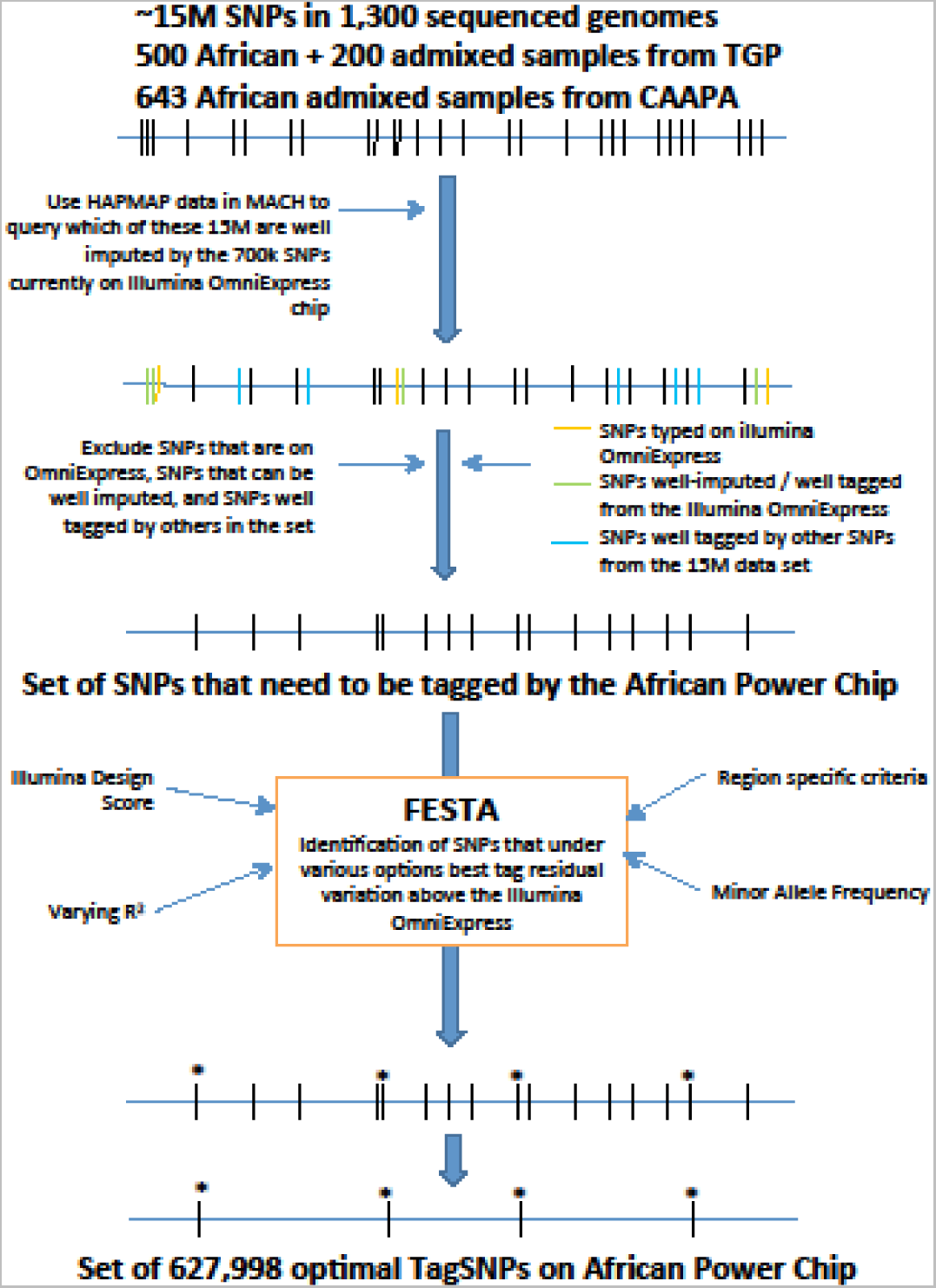
The ADPC design pipeline, describing the steps taken to whittle ~15 million novel African SNPs into a 627k African-targeted GWAS array

## Results

Based on the combination of OmniExpress and the ADPC, coverage is exceptionally good in both the 1000 Genomes African and admixed African populations. [Figure 2] The average r^2^ for all variants is greater than 0.8 at ≥1% MAF. Coverage is slightly better amongst admixed African populations than continental African populations, which is useful for the study of African Americans in particular. It is also not surprising, given that we had only one continental African population, compared to 15 African-admixed populations. In all populations, this represents a much better coverage level than with previous commercially available arrays, and represents an important step forward for studies of individuals of African descent.

**Figure 2.**
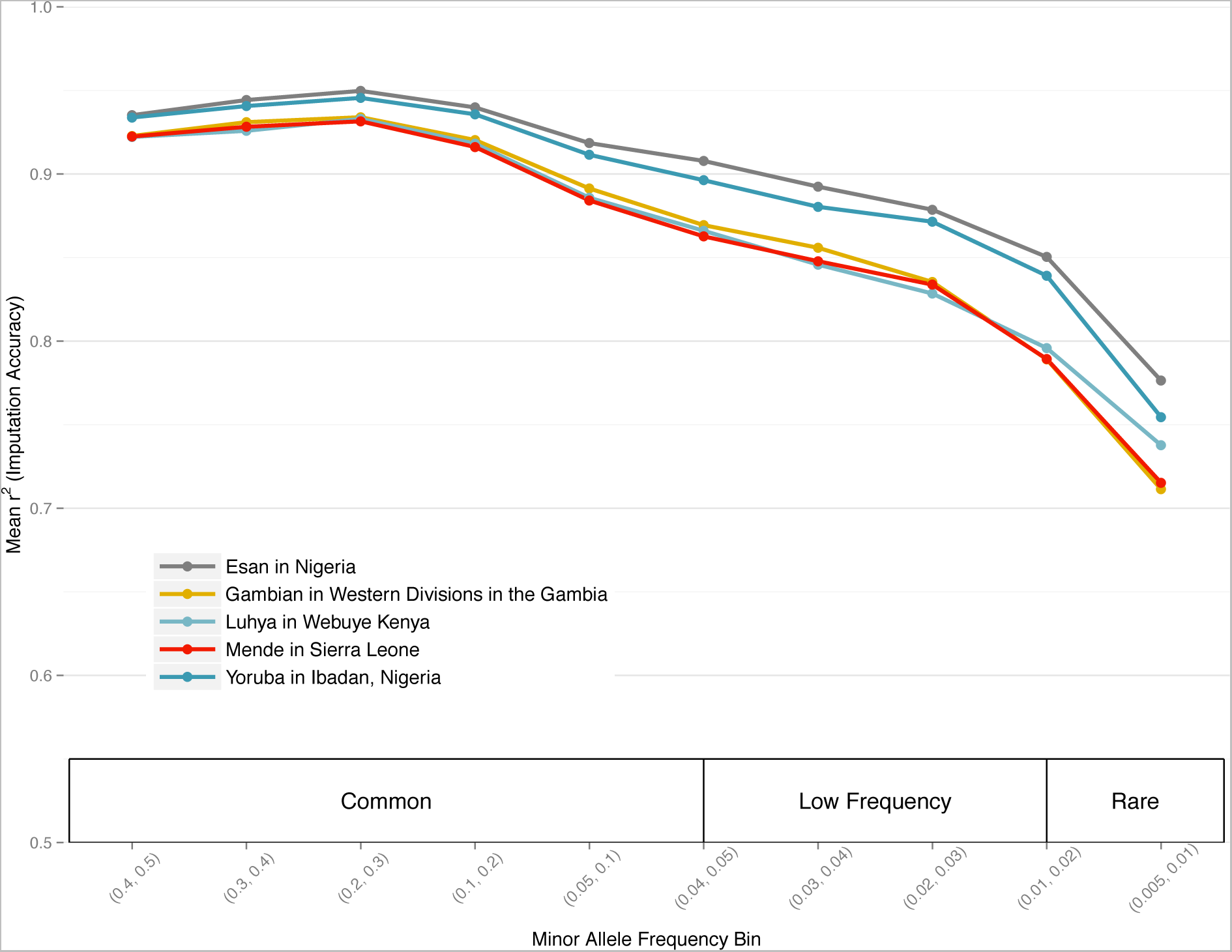
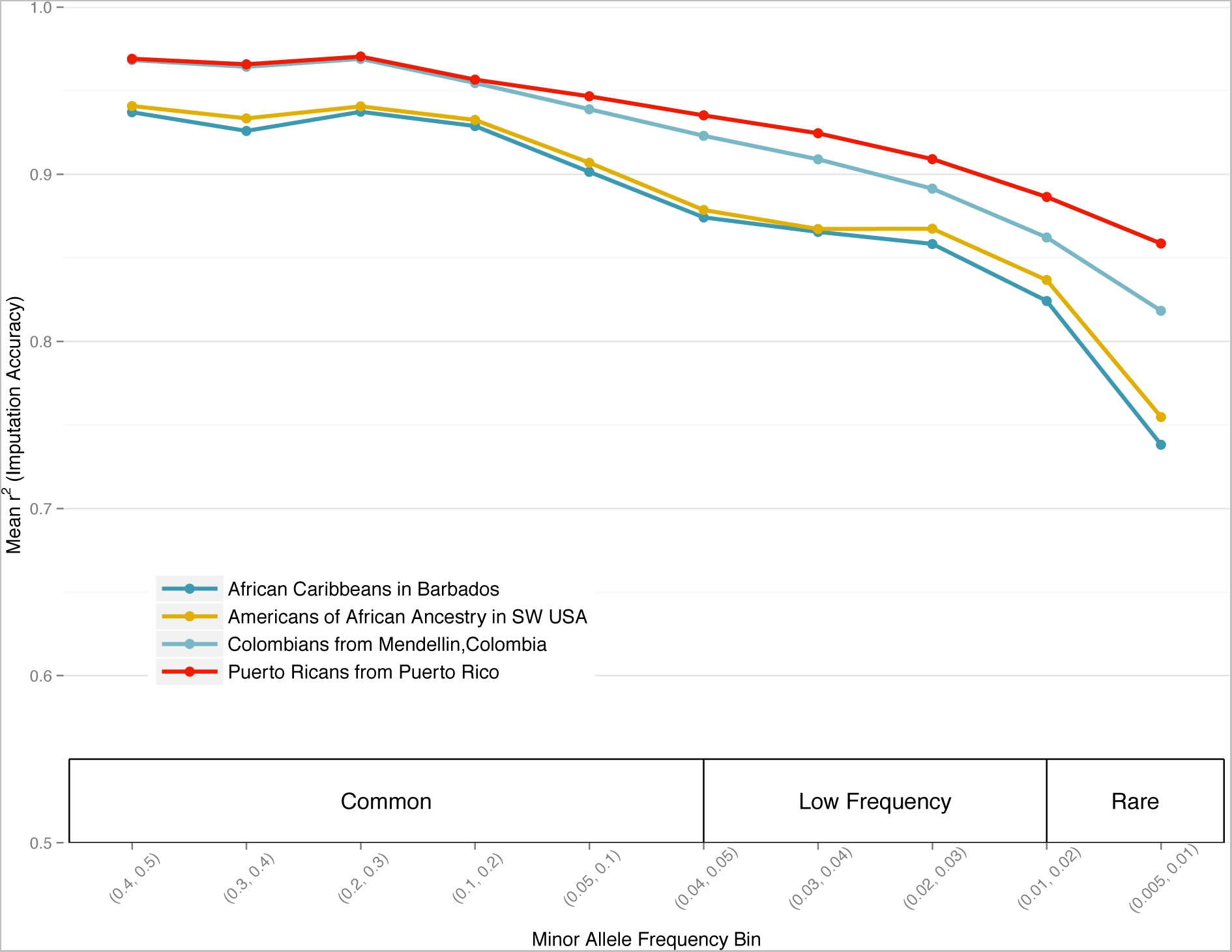
Estimated imputation coverage of variants tagged by the combination of the ADPC and OmniExpress. **2a**. Coverage in 1000 Genomes African populations is >= .8 r^2^ down to 1% MAF **2b.** Coverage in 1000 Genomes admixed African populations is >= .8 r^2^ down to 1% MAF

Additional analysis of SNP coverage at ≥ 1% MAF in the CAAPA population was conducted to ensure our array will enable researchers to have sufficient power to identify novel associations between disease phenotypes and low frequency variants specific to African populations. Despite raising the MAF threshold for TagSNPs to 1.6%, we report coverage of all variants with MAF greater than or equal to 1% to give a full picture of the low frequency coverage the ADPC can provide. Genome-wide, the OmniExpress array is estimated to tag 20% of CAAPA variants at *r*^2^ = 0.9, 26% at *r*^2^ = 0.8 and 39% at *r*^2^ = 0.5. All selected variants of the ADPC, alone, are estimated to tag 12% of known variants at *r*^2^ = 0.9, 16% at *r*^2^ = 0.8, and 31% at *r*^2^ = 0.5. The combination of these two arrays is estimated to tag 29% of all CAAPA variants at *r*^2^ = 0.9, 37% at *r*^2^ = 0.8, and 56% at *r*^2^ = 0.5, an improvement of about 50% more variants tagged over the OmniExpress array across the three thresholds. [Table 2]

**Table 2.**
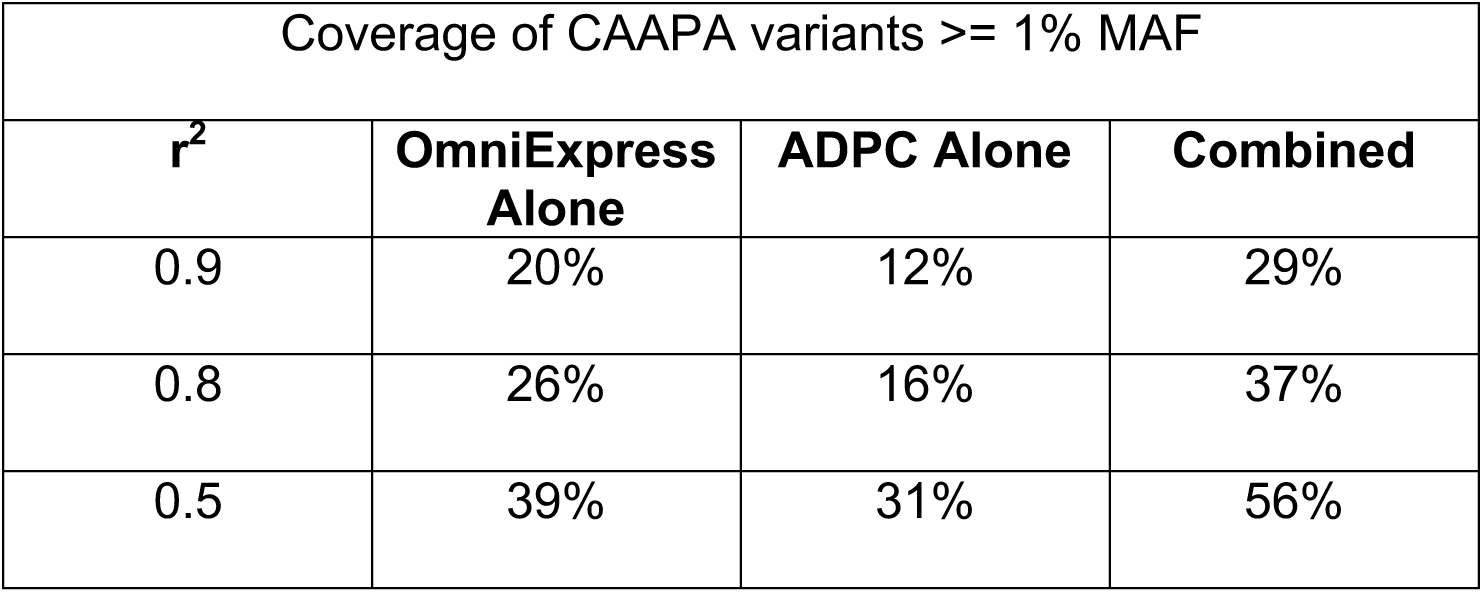
Projected coverage for the ADPC among CAAPA variants >=; 1% MAF, with and without OmniExpress pairing, for the whole genome

While we consider these coverage statistics strong, they only refer to the coverage for variants identified through whole-genome sequencing in CAAPA. This is the most difficult possible test set, since coverage of more common variants in the 1000 Genomes data is not included here. We use this information to give researchers an accurate view of the coverage available for the wealth of novel, low frequency genetic variation identified by CAAPA’s whole-genome sequencing.

Using ~12,000 samples from the CAAPA consortium for the initial run of the ADPC array, ~495,000 out of 700,000 variants passed Illumina’s QC thresholds. This relatively high marker failure rate is not unexpected, however, because the array is comprised of nearly 100% novel markers, never before manufactured. The 494,094 markers successfully manufactured performed excellently, with missing genotype rate averaging only 0.3%.

An important and unique feature of the ADPC is the significantly skewed MAF spectrum of variants on the array [Figure 3]. Compared to OmniExpress, the ADPC contains vastly more low frequency variants. This was not a conscious decision in the design process. Instead, it is the result of trying to tag a set of variants not previously tagged by commercially available genomewide marker arrays. As a result, the combination of the ADPC and OmniExpress is an efficient pairing that increases coverage of the full MAF spectrum for novel variants and dramatically improves the imputation power for low frequency variants.

**Figure 3.**
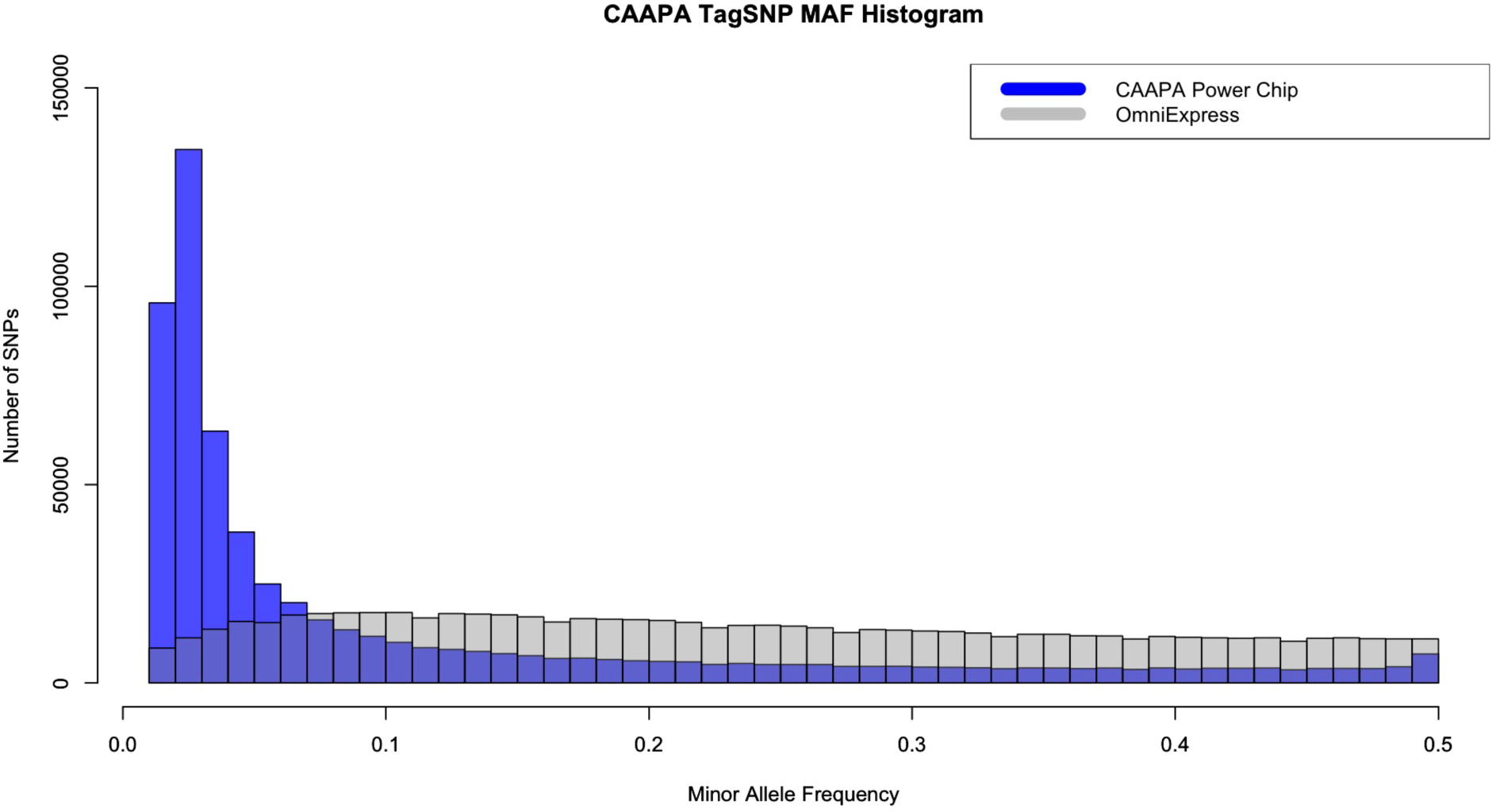
Projected minor allele frequency histograms for the ADPC and OmniExpress arrays overlayed with one another. The disparity between the arrays is significant, and represents very different tagging approaches. This makes them well suited to complement each other.

Through the use of this array, researchers will have greater statistical power to find associations with complex diseases in populations of African ancestry, which has several practical benefits. First, researchers who have already studied populations of African ancestry can inexpensively improve the statistical power of their original studies by adding ADPC genotyping data to their existing GWAS chip data. This chip will provide additional value from its tremendous improvement in the quality of imputed genotypes across the genome as well. At the same time, new researchers will be able to determine power before starting a study in populations of African ancestry, using a combination of the ADPC and OmniExpress, leading to smaller sample sizes needed, and more studies being possible.

## Discussion

In this paper, we present the African Diaspora Power Chip, an affordable genotyping array that dramatically increases the coverage of genetic variants specific to African populations (and their descendants). Through the use of this array, researchers can now be better powered to detect disease associations in populations of African ancestry. We argue that the use of this chip will have a 2-fold benefit: 1) researchers who have already genotyped individuals from populations of African ancestry can inexpensively improve their power of their data by genotyping their subjects on the ADPC, and 2) new studies of African ancestry subjects can now achieve greater power in a study of individuals of African ancestry. For the CAAPA consortium, this means using the ADPC to genotype >13,000 Asthma cases and controls from 9 populations across the Americas.

Although this array was designed to meet the specific needs and timeline of the CAAPA consortium, and may not represent the ideal for a strictly African-based SNP array, it is available to researchers now and has demonstrably increased coverage in variants of West African ancestry. This is, of course, the most common African admixed population in African Americans or other members of the African diaspora, so it is especially well positioned to be useful in studies of admixed African Americans. Furthermore, the CAAPA consortium has released an imputation reference panel, based on the results of our whole-genome sequencing experiment, and it is available through the Michigan Imputation Server (https://imputationserver.sph.umich.edu/index.html), and maximizes coverage provided by the ADPC.

Our immediate plans are to assess the coverage provided by the ADPC in populations of African descent not originating in West Africa. Of specific interest to geneticists are populations in East and North Africa. In the future, through the use of this array, the number of meaningful results from GWAS studies conducted in populations of African descent should increase significantly, providing a more accurate picture of causal disease variants in this group. This will enable personalized medicine techniques to be applied to a much larger subset of Americans than is currently feasible.

## Methods

The 642 individuals in the data freeze were sequenced using Illumina’s Hi-Seq 2000 equipment and the reads were 100bp, paired-end runs. Assembly was performed by the Consensus Assessment of Sequence and Variation (CASAVA) package, which is the Illumina in-house assembly and variant calling technology. The SNP-caller implemented in CASAVA uses a probabilistic model to ultimately generate probability distributions over all diploid genotypes for each site in each genome. A set of MAXGT quality scores is thus generated for each genomic site, corresponding to the ‘consensus quality’ in the SamTools SNP calling method.^14^ These quality scores are then parsed based on a set of consortium-wide rules in order to determine the likely set of variants.

Data processing to generate a 691-sample VCF file for each chromosome from the Illumina MAXGT single-sample SNP VCF files provided in Illumina’s standard deliverable package was performed at Knome, Inc. (Cambridge, MA, USA). The individual VCF files only contained calls for variants, not ref/ref homozygotes. To generate a multi-sample VCF file, these individual VCF files were merged using VCFTools^15^ (v0.1.11), then using custom scripts, a multi-sample VCF file was backfilled to include homozygous reference genotypes and depth of coverage from the sites.txt files. Custom QC scripts confirmed the multi-sample VCFs and the single-sample VCFs had the same number of heterozygous and homozygous alternate genotypes. VCFtools was used to confirm all subjects were included in each multi-sample VCF. The multi-sample VCF was generated including the 48 samples from the SCAALA (Salvador, Brazil) group, but these samples were subsequently dropped from all analyses, leaving 642 individuals, and variants unique to SCAALA were removed from the variant pool. [Mathias et al. 2016]

To pare down the list of variants needing to be tagged, several exclusion sets were created, starting with design score analysis. The segment extending 60 base pairs up and down stream from each variant position were surveyed to determine which side of the variant would create a better probe and a design score was calculated, on a 0-1 scale, representing the estimated success rate for the variant. Any variant scoring below 0.5 was removed. Variants already on the OmniExpress array were also excluded.

To determine which CAAPA variants could be well imputed, the software packages MaCH^9^ and minimac^10^ were employed. All CAAPA samples were first pre-phased by MaCH; subsequently variants still remaining in the tagging pool were imputed in minimac, a low-memory, computationally efficient variant of MaCH, specifically designed for haplotype-to-haplotype imputation. Variants were imputed using the 1000 Genomes Phase I African Reference Panel as the reference.

## Acknowledgments

The authors gratefully acknowledge the contributions of Paul Levett, Anselm Hennis, P. Michele Lashley, Raana Naidu, Malcolm Howitt, and Timothy Roach (**BAGS**), Audrey Grant, Eduardo Viera Ponte, Alvaro A. Cruz, and Edgar Carvalho (**BIAS**), Susan Balcer-Whaley, Maria Stockton-Porter, and Mao Yang (**GRAAD**), Mario Meraz, Jaime Nuñez, Eileen Fabiani Herrera Mejía (**HONDAS**), Deanna Ashley (**JAAS**), Silvia Jimenez, Nathalie Acevedo, Dilia Mercado (**PGCA**), Ann Jedlicka (**REACH**), Addison K. May, Caroline Gilmore, Patricia Minton (**Vanderbilt University**), Qun Niu, (**University of Chicago**), Adeyinka Falusi, Abayomi Odetunde (**University of Ibadan, Nigeria**). The authors also acknowledge the support of John Jay Shannon (**Cook County Health Systems**) and Kevin Weiss (**Northwestern University**), Regina Miranda and the Indians Zenues guards (**San Basilio de Palenque, Bolivar, Colombia**), Ulysse Ateba Ngoa (**Leiden University**), and Charles Rotimi, Adeyemo Adebowale, Floyd J Malveaux, and Elena Reece (**Howard University**). We thank the numerous health care providers and community clinics and co-investigators who assisted in the phenotyping and collection of DNA samples, and the families and patients for generously donating DNA samples to **BAGS, BIAS, BREATHE, CAG, GRAAD, HONDAS, REACH, SAGE II, VALID, SAPPHIRE, SARP, COPDGene, JAAS, GALA II, PGCA,** and **AEGS**. Special thanks to community leaders, teachers, doctors and personnel from health centers at the Garifuna communities for organizing the medical brigades and to the medical students at Universidad Catñlica de Honduras, Campus San Pedro y San Pablo for their participation in the fieldwork related to **HONDAS**; study coordinator Sandra Salazar, and the recruiters in **SAGE and GALA**: Duanny Alva, MD, Gaby Ayala-Rodriguez, Ulysses Burley, Lisa Caine, Elizabeth Castellanos, Jaime Colon, Denise DeJesus, Iliana Flexas, Blanca Lopez, Brenda Lopez, MD, Louis Martos, Vivian Medina, Juana Olivo, Mario Peralta, Esther Pomares, MD, Jihan Quraishi, Johanna Rodriguez, Shahdad Saeedi, Dean Soto, Ana Taveras, Emmanuel Viera, Dr. Michael LeNoir, Dr. Kelley Meade, Mindy Jensen, and Adam Davis; and health liaisons and public health officers of the main Conde office, Adaliudes Conceição, Luciana Quintela, Ivanice Santos, Analú Lima, Benivaldo Valber Oliveira Silva, and Iraci Santos Araujo, and students from the Federal University of Bahia who assisted in data collection in **BIAS**: Rafael Santana, Roberta Barbosa, Ana Paula Santana, Charlton Barros, Marcele Brandão, Ludmila Almeida, Thiago Cardoso and Daniela Costa. We are grateful for the support from the international state governments and universities from Honduras, Colombia, Brazil, Gabon, Nigeria, Netherlands, Jamaica, Barbados and the United States who made this work possible. We also thank Robert Genuario for invaluable assistance in the whole genome sequencing at Illumina, Inc., Gonçalo Abecasis, William Cookson, Miriam Moffatt, for helpful discussions, Pat Oldewurtel and Murali Bopparaju for technical support, Shuai Yuan for software support, and Kit Rees and Cate Kiefe for artistic contributions. We thank Steven Salzberg and Alex Szalay for computing and data storage resources available on the Data-Scope instrument at the Institute for Data Intensive Science (IDIES), Johns Hopkins University. The authors also acknowledge the support from James Kiley, Susan Banks-Schlegel, and Weiniu Gan at the National Heart, Lung, and Blood Institute.

## Funding

Funding for this study was provided by **National Institutes of Health (NIH)** R01HL104608.

Additional NIH funding includes **NCI,** R21CA178706 (RDH), U01CA161032, P50CA125183 (OO). **NCRR**, G12RR003048 (GMD), RR24975 (TH). **NHGRI,** R01HG007644, R21HG007233 (RDH), R21HG004751 (HRJ, JG, ZSQ), T32HG000044 (CRG). **NHLBI,** R01HL087699 (KCB), R01HL118267 (LKW), R01HL117004, R01HL088133, R01HL004464 (EGB), HL081332, HL112656 (LBW), R01HL69167, U01HL109164 (EB,DM), RC2HL101651, RC2HL101543, U01HL49596, R01HL072414 (CO), R01HL089897, R01HL089856, K01HL092601(MGF), R01HL51492, R01HL/AI67905 (JGF), HHSN268201300046C, HHSN268201300047C, HHSN268201300048C, HHSN268201300049C, HHSN268201300050C (JGW). **NIAID,** K08AI01582 (TH), R01AI079139 (LKW), U19AI095230 (CO). **NIEHS**, R01ES015794 (EGB). **NIGMS**, S06GM08016 (MUF), T32GM07175 (CRG). **NIMHD**, P60MD006902 (EGB), 8U54MD007588, P20MD0066881 (MGF). **NSFGRF** #1144247 (RT).

Additional sources of funding include: American Asthma Foundation (LKW and EGB), American Lung Association Clinical Research Grant (TH), Colombian Government (Colciencias) 331-2004 and 680-2009 (LC), EDCTP:CT.2011.40200.025 (AAA), EU-IDEA HEALTH-F3-2009-241642 and EU-TheSchistoVac HEALTH-Fe-2009-242107 (MY), Ernest Bazley Fund (PCA, RK, LG, RS), and the Fund for Henry Ford Hospital (LKW). The Jamaica 1986 Birth Cohort Study was supported by grants from the Caribbean Health Research Council, Caribbean Cardiac Society, National Health Fund (Jamaica) and Culture Health Arts Sports and Education Fund (Jamaica). Study nurses were supported by the University Hospital of the West Indies (TF,JKM), Ralph and Marion Falk Medical Trust (COO,OO,OO,GA), UCSF Dissertation Year Fellowship (CRG), Universidad Católica de Honduras, San Pedro Sula (EHP), University of Cartagena (JM), Wellcome Trust 072405/Z/03/Z, 088862/Z/09/Z (PJC). The Jackson Heart Study is supported by contracts HHSN268201300046C, HHSN268201300047C, HHSN268201300048C, HHSN268201300049C, HHSN268201300050C from the NHLBI and the NIMHD. EGB was funded by Flight Attendant Medical Research Institute, RWJF Amos Medical Faculty Development Award and the Sandler Foundation; the Sloan Foundation to RDH; CRG was supported in part by the UCSF Chancellor’s Research Fellowship and Dissertation Year Fellowship.

KCB was supported in part by the Mary Beryl Patch Turnbull Scholar Program. RAM was supported in part by the MOSAIC Initiative Awards from Johns Hopkins University. MP-Y was funded by a Postdoctoral Fellowship from Fundación Ramón Areces. MIA is an investigator supported by National Council for Scientific and Technological Development (CNPq). TVH was supported in part by K24 AI 77930, UL1 TR00445, and U19 AI95227. RO was funded by NHLBI Diversity Supplement R01HL104608.

Funding for the cohorts was provided by the following: AEGS, BAGS, BIAS, BREATHE (K08AI001582, RR24975), CAG, COPDGene, GALA II, GRAAD, HONDAS, JAAS(The Jamaica 1986 Birth Cohort Study was supported by grants from the Caribbean Health Research Council, Caribbean Cardiac Society, National Health Fund (Jamaica) and Culture Health Arts Sports and Education Fund (Jamaica). The study nurses were supported by the University Hospital of the West Indies.), PGCA (University of Cartagena and Colciencias Contracts 183-2002, 680-2009), REACH, SAGE II, SAPPHIRE, SARP, SCAALA, VALID.

Individual-level sequence data presented in this study are currently being prepared for deposition into dbGAP. Currently, these data are available through a data access agreement to respect the privacy of the participants for the transfer of genetic data, by contacting R.A.M. and K.C.B. and CAAPA (http://www.caapaproject.net).

## Author contributions statement

HRJ designed and performed experiments, analyzed data and wrote the manuscript. JG, YJH, ZSQ, RAM, THB and KCB conceived the experiment and critically reviewed the manuscript. MH, DB, RG and CL provided technical assistance. PAG and AL provided HLA expertise. TO, GA, EK, GW, CG and CB performed experiments, generated figures and assisted with the writing and review of the manuscript.

## Accession codes for deposited data

The whole genome sequence data that support the findings of this study have been deposited in dbGAP with the accession code phs001123.v1.p1.

## Competing financial interests

Mark Hansen, Rob Genuario, Dave Bullis and Cindy Lawley are (or were) Illumina employees.

